# Early and widespread cerebellar engagement during hippocampal seizures and interictal discharges

**DOI:** 10.1101/2024.05.14.593969

**Authors:** M.L. Streng, B.W. Kottke, E.M. Wasserman, L. Zecker, L. Luong, S. Kodandaramaiah, T.J. Ebner, E. Krook-Magnuson

**Affiliations:** Department of Neuroscience, University of Minnesota, Minneapolis, MN, USA; Department of Mechanical Engineering, University of Minnesota, Minneapolis, MN, USA

**Keywords:** Temporal Lobe Epilepsy, See-Shell, Purkinje cells, IED, focal epilepsy

## Abstract

Despite research illustrating the cerebellum may be a critical circuit element in processes beyond motor control, and growing evidence for a role of the cerebellum in a range of neurological disorders, including the epilepsies, remarkably little is known about cerebellar engagement during seizures. We therefore implemented a novel method for repeated widefield calcium imaging of the cerebellum in awake, chronically epileptic mice. We found widespread changes in cerebellar Purkinje cell activity during temporal lobe seizures. Changes were noted in the anterior and posterior cerebellum (lobules IV-VII), along the midline (vermis), and both ipsilaterally and contralaterally (in the simplex and Crus I) to the seizure focus. This was true for both overtly behavioral seizures *and* for hippocampal seizures that remained electrographic only -- arguing against cerebellar modulation simply reflecting motor components. Moreover, even brief interictal spikes produced widespread alterations in cerebellar activity. Perhaps most remarkably, changes in the cerebellum also occurred *prior* to any noticeable change in the hippocampal electrographic recordings. Together these results underscore the relevance of the cerebellum with respect to seizure networks, warranting a more consistent consideration of the cerebellum in epilepsy.

## Introduction

The cerebellum is canonically associated with motor learning, but it is increasingly appreciated for its role in a wide range of physiological processes ^1–9^. It is estimated that the human cerebellum contains more than half of all the neurons in the entire central nervous system ^10^, and across species has ∼3.6 times as many neurons as the cerebral cortex ^11^. The cerebellum is functionally connected, often via reciprocal loops, with a wide range of brain regions ^12,13^.

Multi-synaptic functional connectivity between the cerebellum and the neocortex is extremely well documented ^14–18^, and there is a growing body of research illustrating such functional connectivity also with allocortex, including the hippocampus ^19–21^. Similarly, while the cerebellum’s role in certain motor disorders has long been recognized ^22–26^, the cerebellum is increasingly implicated in a wider range of neurological disorders ^27–29^, including the epilepsies^30,31^.

While the cerebellum is often neglected in epilepsy research and clinical care alike, the cerebellum can be impacted by epilepsy and seizures ^30–37^, and vice versa ^38–42^. Interventions targeting the cerebellum can inhibit a range of seizure types ^30,31,36,40–43^, including but not limited to hippocampal seizures ^33,38,39,44^. While certain anti-seizure medications can directly impact the cerebellum, epilepsy-related cerebellar deficits extend beyond these impacts ^30,45,46^, and can even be seen in animal models ^47,48^. This indicates that epilepsy itself is impacting the cerebellum. Changes in cerebellar gray matter in epilepsy patients is associated with future SUDEP (sudden unexpected death in epilepsy) events ^45^. The cerebellum may therefore have a particularly significant role in epilepsy and epilepsy outcomes. In addition to the impacts of chronic epilepsy on the cerebellum, research indicates that the cerebellum itself can seize (recently reviewed in reference ^30^).

Unfortunately, even in rodents, studies examining cerebellar engagement during seizures have been relatively limited ^30,32–34^. Depth EEG and LFP recordings have shown engagement of the cerebellum during seizures ^30^, including in human patients ^34,49,50^. Single unit recordings have additionally overcome potential volume-conduction concerns associated with field recordings, illustrating that seizures can modulate the activity of individual Purkinje cells ^32,33,36,51^. While these studies have convincingly demonstrated that the cerebellum can “seize” ^32,33,52^, the extent of such modulation is unknown, as is the consistency of such responses across multiple spontaneous seizure events. Previous work using calcium imaging has yielded important insights into seizures and other epileptic networks ^53–73^, and provides an opportunity to study the cerebellum in epilepsy. Specifically, a method to allow repeated, widefield, calcium imaging of the cerebellum in chronically epileptic animals could allow examination of activity across a large area of the cerebellum during multiple spontaneous epileptiform events.

We therefore adapted conformant See-Shell ^74^ technology to allow repeated widefield monitoring of calcium activity in Purkinje cells across the dorsal cerebellar surface, including vermal lobules IV-VII as well as portions of the simplex and Crus I/II ^75^ in chronically epileptic animals. Concurrent hippocampal LFP recordings allowed for monitoring of epileptiform activity. Sparse expression of GCaMP6s selectively in Purkinje cells allowed single cell resolution, and 1-photon imaging allowed simultaneous monitoring of both somatic and dendritic calcium signals ^75^. Repeated imaging sessions in chronically epileptic animals allowed us to capture even relatively rare, spontaneous, large, overtly behavioral, seizure events -- in addition to more frequent electrographic hippocampal seizure events and interictal epileptiform discharges (IEDs).

Overtly behavioral seizures produced *widespread* changes in the cerebellum, including in the anterior and posterior cerebellum, along the midline, and both ipsilaterally and contralaterally to the seizure focus, often beginning several seconds before any observed electrographic or behavioral changes. Notably, hippocampal seizures that remained electrographic only *also* produced widespread cerebellar changes -- arguing against cerebellar modulation simply reflecting motor components. Even brief IEDs were associated with widespread alterations in cerebellar activity, indicating also that sustained hippocampal alterations were not required to produce cerebellar changes. Perhaps most remarkably, across all three forms of epileptiform activity, changes in the cerebellum could occur *prior* to any noticeable change in the hippocampal LFP recordings.

These findings highlight the functional connectivity between the hippocampus and the cerebellum, and illustrate the potential consequences of this functional connectivity in neurological disorders. Specifically, even a disorder believed to have its focus in the hippocampus can have large impacts on the cerebellum. Cerebellar changes may even be detectable prior to detectable changes in the hippocampus. Together, these findings underscore that the cerebellum is an extremely relevant brain structure in epilepsy, and warrants more consistent consideration in epilepsy research and care.

## Methods

### Ethical approval

All experimental protocols were approved by the University of Minnesota’s Institutional Animal Care and Use Committee.

#### Animals

Mice were bred in-house and had *ad libitum* access to food and water in all housing conditions. Mice expressing the Ca^2+^ indicator GCaMP6s selectively in Purkinje cells were generated by crossing mice expressing Cre selectively in Purkinje cells in the cerebellum (B6.Cg-Tg(Pcp2-cre)3555Jdhu/J; Jackson Laboratory stock 010536)^76^ with floxed-STOP GCaMP6s mice (B6.Cg-Igs7tm162.1(tetO-GCaMP6s,CAG-tTA2)Hze/J; Jackson Laboratory stock 031562)^77^. The resulting PcP2-GCaMP6s crosses had expression of GCaMP6s restricted to Purkinje cells, and in a sparse manner, as reported previously^75^. This allowed for single cell resolution of Purkinje cells at the wide-field level. Until implantation, animals were housed in standard housing conditions in the animal facility at the University of Minnesota. Following implantation, animals were singly housed in investigator managed housing on a 12-hour reverse light/dark cycle. A total of n = 6 mice of both sexes (n = 3 M, n = 3 F) were utilized in this study.

### Surgical procedures

#### Epilepsy induction

Procedures for epilepsy induction using the mouse unilateral intrahippocampal kainic acid model of Temporal Lobe Epilepsy (TLE) largely followed previously published protocols ^39,44,78–81^. Specifically, animals postnatal day 45 or greater were injected with 100nL kainic acid (KA, Tocris Biosciences) unilaterally into the right dorsal hippocampus (2.0 mm posterior, 1.25 mm right, 1.6 mm ventral from bregma) under isoflurane anesthesia (1.5-2%) as done in our previous work^39,44^. Animals were removed from isoflurane within five minutes post injection^82^. In this model, spontaneous recurrent electrographic seizures emerge from the damaged hippocampus, providing a strong model of pharmocoresistant ^83,84^ TLE with hippocampal sclerosis ^79,83,85,86^. Post-operative care consisted of recovery from anesthesia on a heating pad with regular visual inspection, followed by daily post-operative monitoring for a minimum of three days to inspect comfort level and healing of the surgical site. Neopredef powder was applied to the closed incisions as a topical antibiotic and analgesic. Only animals that showed spontaneous electrographic hippocampal seizures weeks after kainate injection were included for simultaneous cerebellar imaging and hippocampal LFP monitoring.

#### Implantation

Cerebellar imaging windows were fabricated as described previously^75^. Implants consist of a small section of PET film affixed to a 3D-printed window frame, designed to conform to the contours of the skull, and a separate 3D-printed head fixation implant, which contained three holes tapped using an 0-80 hand tap for securing to a titanium headplate during the implantation surgery. A minimum of 2 weeks post kainate injection, mice underwent implantation procedures^75^ under isoflurane anesthesia. Buprenorphine ER (1mg/kg) and Dexamethasone (2mg/kg) were administered subcutaneously for analgesia and inflammation, respectively. The scalp was removed to expose the skull over the dorsal surface of both the cerebral and cerebellar cortices, and a large craniotomy matching the external profile of the cerebellar window frame was performed, exposing lobules IV/V, VI, and VII of the cerebellar cortex. The window was placed over the cerebellar cortex and bonded to the skull using Metabond dental cement (C&B Metabond, Parkell Inc.). Following window implantation, a twisted wire bipolar (local reference, differential) electrode (Protech International) was stereotactically implanted in the hippocampus, ipsilateral to the site of kainate injection, at 2.6 mm posterior, 1.75 mm right, 1.6 mm ventral from bregma, and secured with Metabond. The head fixation implant was next placed on the skull over the dorsal cerebral cortex and secured using Metabond. A custom titanium headplate was screwed onto the previously tapped holes in the head fixation implant, and the entire space between the window and headplate was filled with dental cement to protect the window from the external environment. A 3D-printed cap was attached to the titanium headplate to protect the PET surface. Post-operative care consisted of recovery from anesthesia on a heating pad with regular visual inspection, followed by daily post-operative monitoring for a minimum of three days to assess comfort level and healing of the surgical site. Animals were allowed to recover a minimum of five days prior to simultaneous cerebellar imaging and hippocampal local field potential (LFP) monitoring.

### Simultaneous fluorescence imaging and hippocampal LFP monitoring

During imaging sessions, mice were head fixed on a freely moving disk treadmill to allow for spontaneous locomotion (Figure 1). Awake, mesoscale cerebellar imaging was performed under an epifluorescence microscope (Nikon AZ100) with a 2X objective, as described previously^75^.

**Figure 1.**
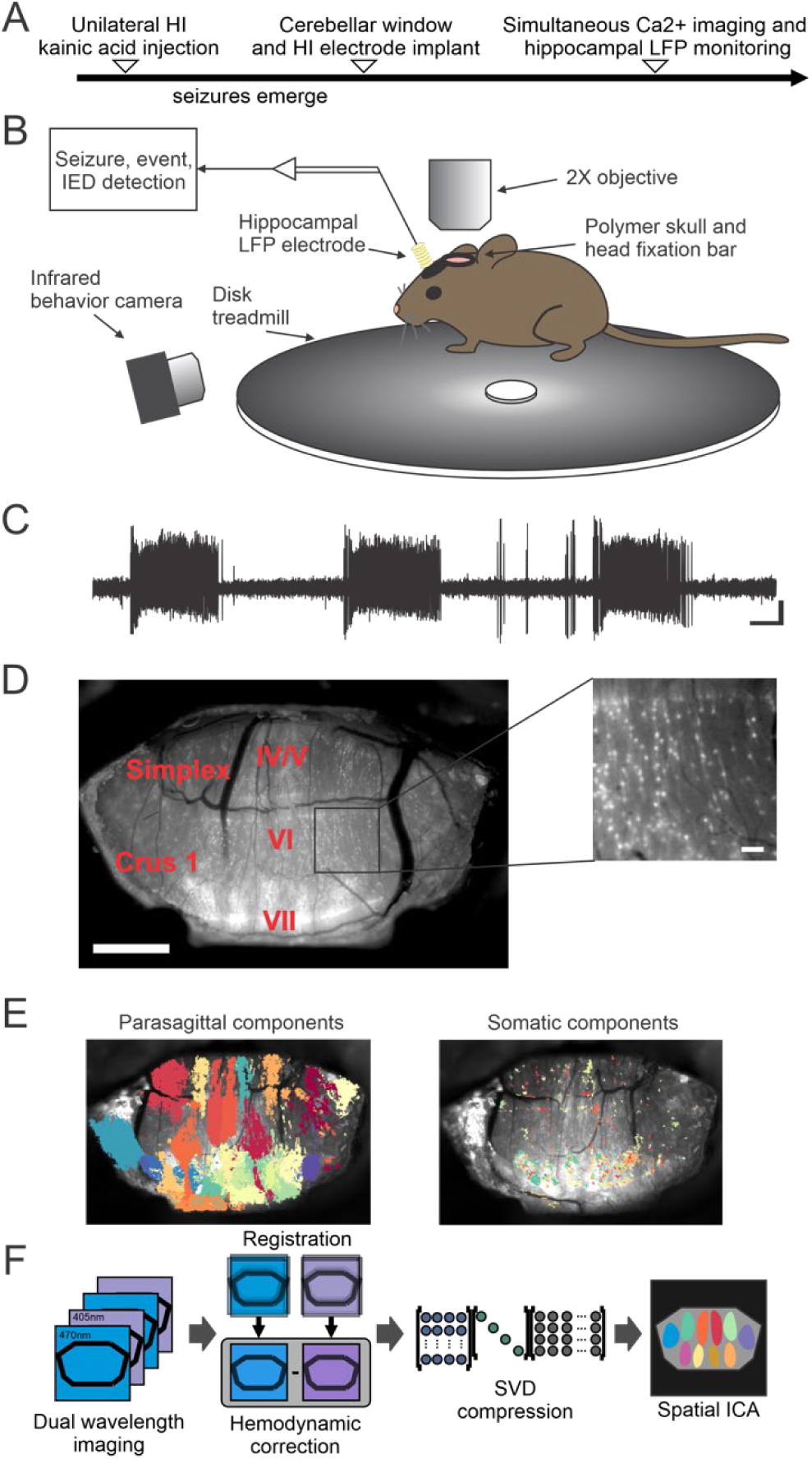
Repeated widefield cerebellar calcium imaging with concurrent hippocampal local field potential recordings in awake, chronically epileptic, animals. A. Experimental timeline. Animals were first injected with kainate into the dorsal hippocampus, to induce temporal lobe epilepsy (TLE). Animals were then implanted with an electrode into the dorsal hippocampus near the original site of kainate injection, to record spontaneous hippocampal epileptiform activity during the chronic phase of the disorder. Animals were additionally implanted with a cranial window over the cerebellum using a modification of See-Shell technology, to allow for concurrent, longitudinal, cerebellar imaging. B. Schematic of recording setup. During imaging sessions, the animal was headfixed on a freely spinning disk treadmill, the hippocampal LFP was recorded, and aligned to the widefield cerebellar imaging. C. Example hippocampal LFP trace, illustrating spontaneous epileptiform activity. Scale bars: 20s, 0.2mV. D. The cerebellar imaging window allowed simultaneous one-photon (1P) imaging of a large area of the cerebellar dorsal cortex, including Vermis lobules IV-VII, Simplex, and Crus I. Scale bar: 1mm. Inset: sparse GCaMP6s expression in Purkinje cells allowed for single cell resolution. Scale bar: 100μm. E. Spatial independent component analysis (sICA), a blind source separation algorithm, resulted in automatic parsing of parasagittal (left) and somatic (right) calcium signals. F. Visual summary of the preprocessing pipeline for wide-field cerebellar imaging data and resulting independent components (ICs). Interleaved dual wavelength imaging (470nm for calcium-dependent signals, 405nm for isosbestic control) and registration of acquired images allowed for correction of movement and hemodynamic changes. Following singular value decomposition (SVD) compression, regions of interest were generated using sICA.

Using a digital zoom, the field of view was adjusted to image the dorsal cerebellum, with the focal plane adjusted to allow for resolution of both Purkinje cell dendrites and somata. Dual wavelength illumination was used to capture both calcium-dependent (470 nm, blue light) and calcium-independent (405 nm, violet light) GCaMP6s signals on consecutive frames by a Cairn OptoLED driver (Cairn OptoLED, P1110/002/000; P1105/405/LED, P1105/470/LED). Calcium-dependent and calcium-independent signals were sampled by alternating the illumination of the LEDs, as described previously^75,87–89^. Single-photon fluorescence images were captured at a frequency of 20 frames per second, 18 ms exposure, using a high-speed CMOS camera (ORCA-Flash4.0 V3, Hamamatsu photonics) controlled with MetaMorph. Behavior was monitored and recorded using a high-speed IR-sensitive CMOS camera (Blackfly, FLIR systems) under infrared illumination that did not affect concurrent calcium imaging.

Synchronization of the behavior and microscope cameras was controlled by a series of TTL pulses delivered by Spike2 software and a CED power 1401 data acquisition system (Cambridge Electronic Design). Hippocampal LFP was recorded via electrical patch cords through an electrical commutator (PlasticsOne), amplified (Brownlee) and digitized (National Instruments). The TTL pulses controlling the behavior and microscope cameras were simultaneously recorded, allowing for the alignment of the Hippocampal LFP data to the cerebellar imaging data. The timing of spontaneous behavioral seizures, electrographic events, and interictal spikes were determined offline using our custom in-house software, as described previously ^38,39,44,86,90^. Spontaneous overt, behavioral seizures were confirmed using the behavioral monitoring videos. Mouse forepaw position was labeled using DeepLabCut for markerless behavioral tracking^91^ to determine movement relative to hippocampal epileptiform activity.

### Spatial independent component analysis

The preprocessing of cerebellar imaging data and determination of Purkinje cell spatial independent components (ICs) largely follows our previous methods^75,88,92–94^. Briefly, the first 10 seconds of each imaging stack data were removed to eliminate the initial rundown of the calcium fluorescence signal. Calcium-independent signals were removed by deinterleaving the 470 nm and 405 nm frames and subtracting the 405 nm signal, as described previously ^87–89,92,95,96^, followed by standard motion correction using co-registration ^75,88^ (**Fig. 1D**). The ΔF/F_avg_ (listed simply as of ΔF/F throughout) for each pixel was computed, with F_avg_ representing the mean corrected calcium-dependent signal across the recording stack. The resulting preprocessed imaging stacks were concatenated for a given recording day. Concatenated stacks were compressed using singular value decomposition (SVD), retaining the first 1000 principal components, which maintains important features of the full-rank data while drastically reducing the necessary computing time ^94,97^. Spatial ICA was performed on the compressed dataset, and significant Purkinje cell ICs were determined using thresholding, and visually inspected for artifacts, as described previously^75^. As observed previously^75^, cerebellar signals consisted of either parasagittal ^98–102^ or somatic domains^75^ (**Fig. 1E**).

### Analysis of cerebellar modulation during epileptiform activity

For recording days with spontaneous overt, behavioral seizures, we aligned the ΔF/F of each cerebellar IC to the onset of large amplitude spiking (i.e. the first large amplitude LFP spike of the seizure event) and offset (defined as the last large amplitude LFP spike prior to onset of post-ictal depression) to determine the timing of cerebellar modulation. For visualization purposes of parasagittal ICs, these ΔF/F signals were converted into a heat map for each parasagittal IC, and displayed at the appropriate spatial location on a map of the imaging window (as shown e.g. in **Fig. 2A, top**). For visualization purposes of somatic ICs, representative somatic ΔF/F signals were plotted as colored traces, with the approximate location of each displayed soma represented by the color of the trace and marked in the accompanying schematic (as shown e.g. in **Fig. 2A, bottom**). For additional visualization of cerebellar calcium signals during individual seizure events, the ΔF/F of each IC over time was separated by the side relative to the seizure focus (i.e. the KA injected hippocampus), averaged into 50 micron bins across the anterior-posterior extent of the window based on the IC center, and color-coded based on the anterior-posterior positioning of the IC center (illustrated e.g. in **Fig. 3A**). In order to determine significant IC modulation with each behavioral seizure, the ΔF/F signal for each IC was aligned to the onset of large amplitude spiking, and was transformed into a z score based on the average interictal activity for that same IC. IC modulation greater than ±3 SD was considered significant. For visualization purposes, this significance threshold was used to create thresholded heatmaps of significant modulation for example seizures by IC location (as illustrated e.g. in **Fig.3A, bottom**). The first time point within 10s of the onset of high amplitude spiking showing a significant modulation was computed for all ICs with significant modulation to determine the relative timing of significant modulation (presented **in Fig 3D-F**). Timing of significant modulation relative to seizure onset versus IC anterior-posterior location was then examined using Pearsons’ correlation. To visualize the two-dimensional timing and magnitude of IC modulation, the timing of modulation onset and peak change in activity for each significantly modulated IC were partitioned based on the normalized IC center into a 10×10 grid, and averaged (as shown in **Fig. 3F**).

**Figure 2.**
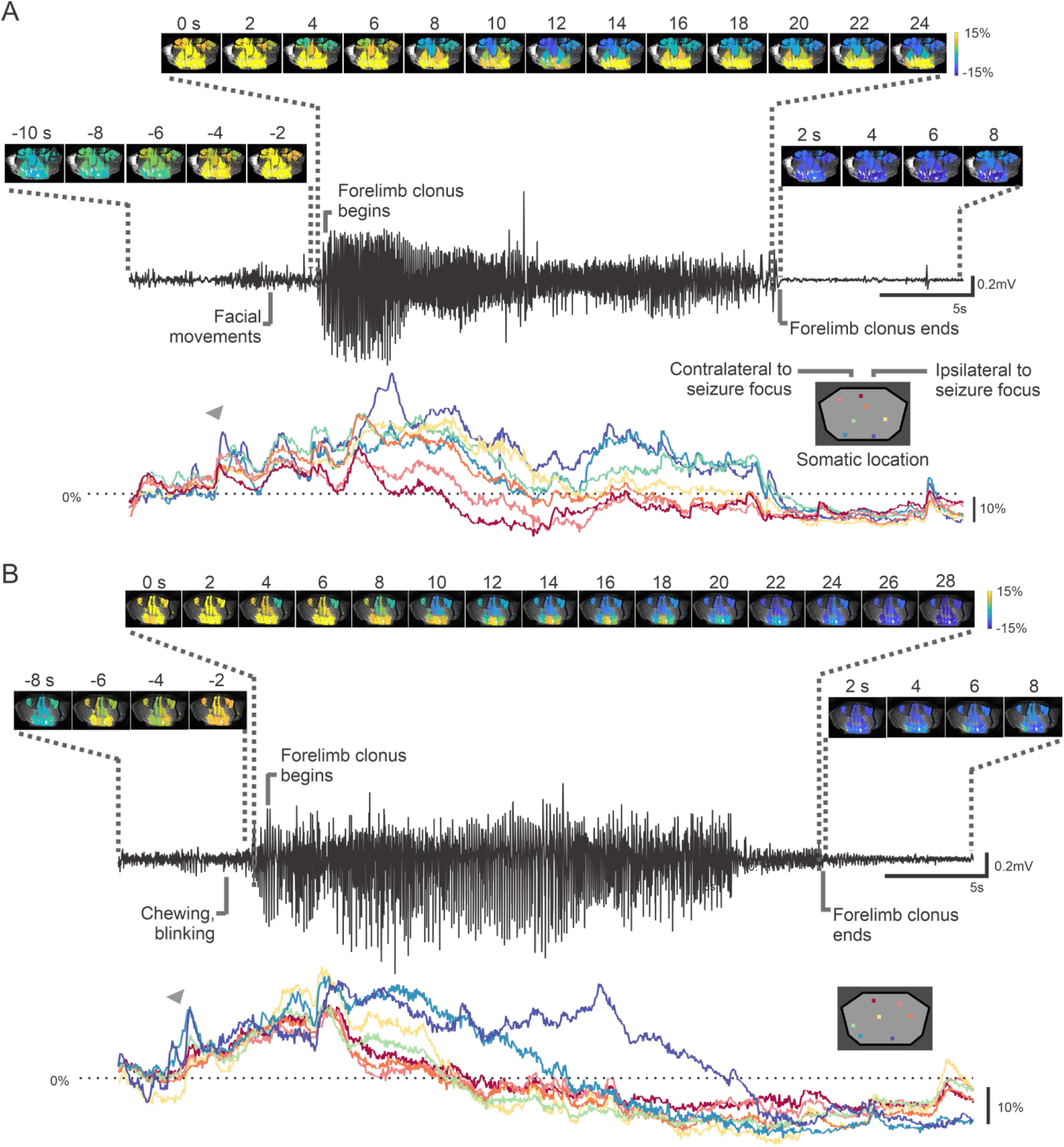
Widespread and early changes in cerebellar activity associated with overtly behavioral seizure events. A. Example recorded spontaneous behavioral seizure (LFP recorded from the dorsal hippocampus, black trace) and associated changes in cerebellar parasagittal domains’ calcium signals (spatial heat maps, top), and ΔF/F traces for example Purkinje cell somata, color coded by location (bottom). Note the widespread changes, including in the contralateral (left in spatial maps) and ipsilateral (right in spatial maps) cerebellum. Note also the increase (gray arrow) that happens prior to motor manifestations, and the widespread decrease during hippocampal postictal depression. B. As in A, but for a different example seizure from a different example mouse. Note again the early increase in cerebellar calcium activity (gray arrow; prior to any noticeable behavioral changes), and the late-ictal decrease (beginning prior to ictal offset), extending into and including strong depression during hippocampal-LFP postictal depression.

**Figure 3.**
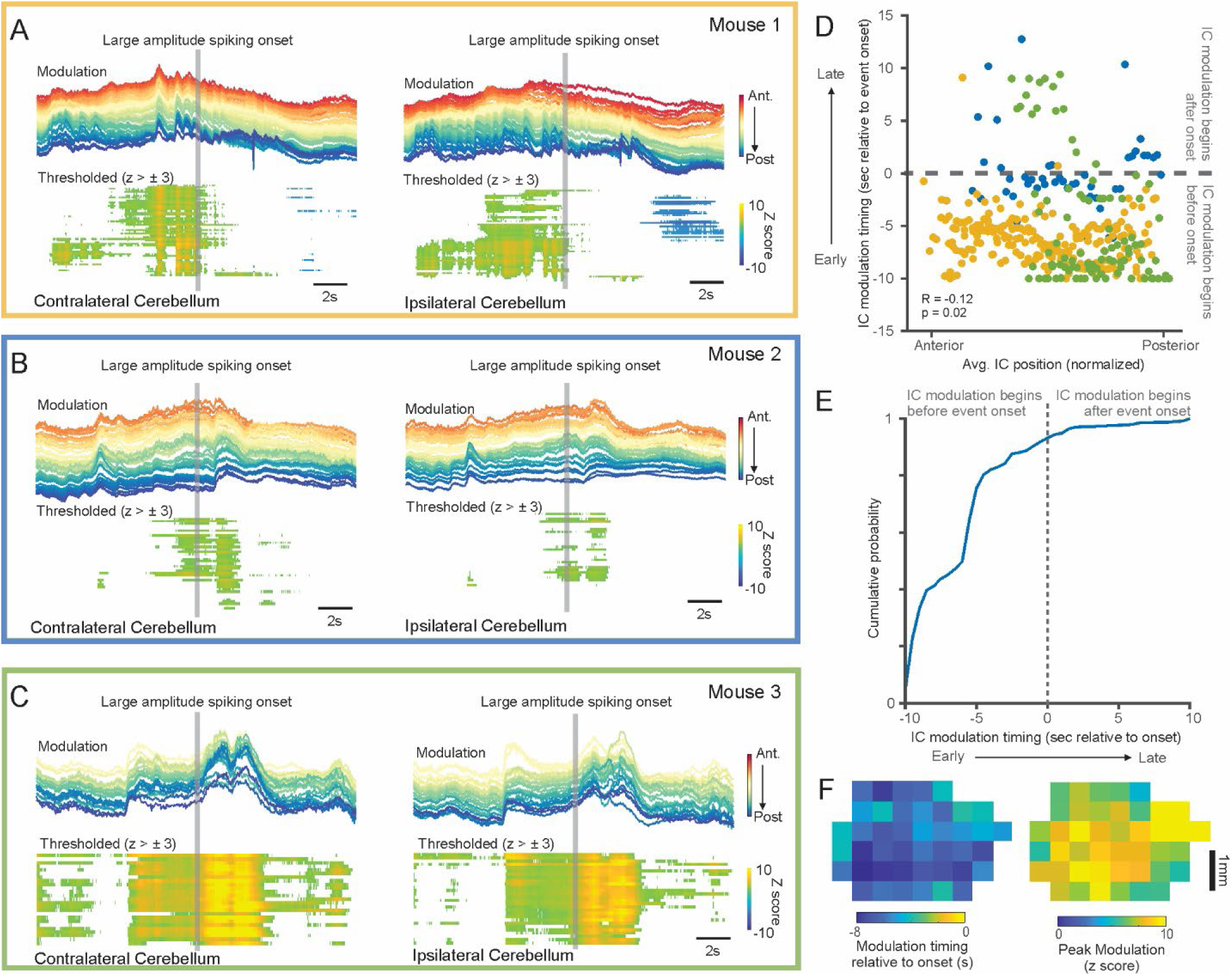
Early cerebellar changes occur across animals and occur broadly across the cerebellum. A. Cerebellar calcium changes for an example large, overtly behavioral, seizure in an example animal. Rainbow plots (top) are the average change in fluorescence across ICs contralateral (left) and ipsilateral (right) to the site of KA injection, with warmer colors denoting anterior cerebellar ICs and cooler colors posterior cerebellar ICs. Bottom heat plots are significance-thresholded activity, illustrating significant increases in cerebellar activation beginning several seconds prior to onset of high amplitude spiking in the hippocampal LFP. B-C Same as A, but for two other example mice with recorded spontaneous, overtly behavioral, seizures. D. Average timing of significant IC modulation relative to the onset of large amplitude spiking based on IC Anterior-Posterior positioning, color coded by mouse. Note that the majority show modulation prior to onset of high amplitude hippocampal ictal spiking. Data drawn from n=10 overtly behavioral seizures in which the onset of the seizure was captured during imaging. E. Cumulative probability distributions for all significantly modulated ICs, indicating significant alterations in IC activation begin before spiking onset across the majority of ICs. F. Averaged timing and peak modulation across all ICs mapped to the cerebellar window, illustrating broad, bilateral increases in cerebellar activity beginning prior to onset of high amplitude ictal spiking in the hippocampal LFP.

**Figure 4.**
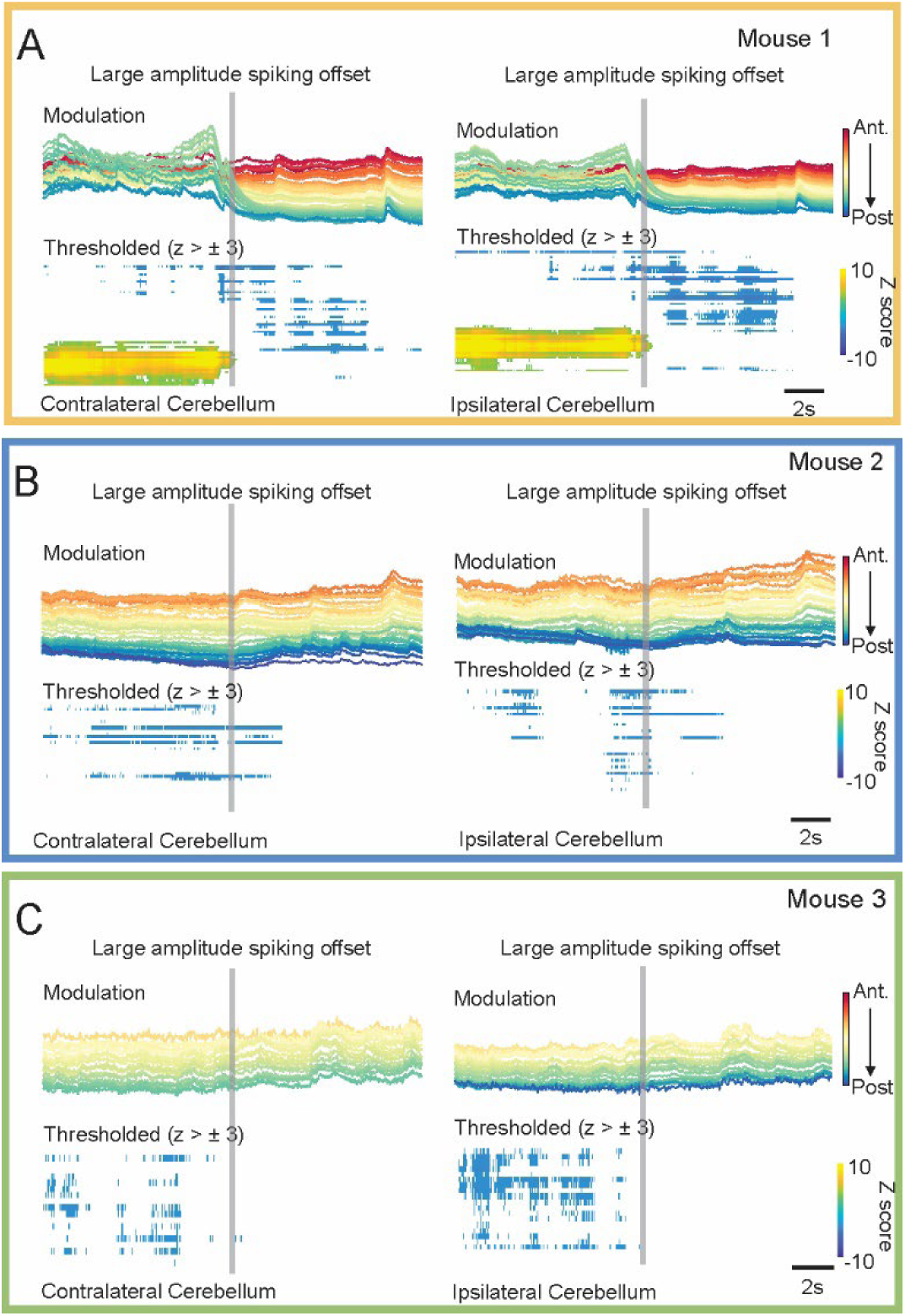
Cerebellar activity is depressed at or prior to the end of behavioral seizures. A. Cerebellar activity aligned to the offset of large amplitude spiking during an example large behavioral seizure captured during wide-field imaging in an example mouse. Conventions are as in Figure 3. B-C. Same as A, but for two other example mice. Note that while some ICs for some seizure events remain elevated until the onset of postictal depression (posterior ICs in examples in A), many show broad decreases prior to seizure offset (examples in B and C). None show continued elevation in the postictal period.

For hippocampal electrographic-only seizure events, the ΔF/F of each cerebellar IC was aligned to the onset or offset of electrographic events (defined as the first and last ictal spike, respectively). Mean population changes in cerebellar activation relative to electrographic event onset and offset were computed by averaging across all ICs for each individual event, as in the heat maps **Fig. 5A & F**, as well as the overall population average across all events to generate an overall mean trace as in **Fig. 5A & F bottom traces**. To examine cerebellar changes ipsilaterally and contralaterally along the anterior-posterior axis, ICs signals were binned by anterior-posterior location, averaged using the same bins as for behavioral seizures, and displayed color-coded by anterior-posterior location (as illustrated e.g. in **Fig. 5C**). In order to determine the significance of IC modulation relative to event onset and offset, the averaged ΔF/F was transformed into a z score based on a shuffled distribution of the same number of time periods as electrographic events, chosen at random from interictal time periods, repeated 1000 times. The timing of decreases and increases relative to event onset and offset, respectively, were determined for each significantly modulated IC, by taking the first time point at which the IC’s activity crossed a 2SD threshold. Spatial timing and peak modulation maps were computed as for behavioral seizures.

**Figure 5.**
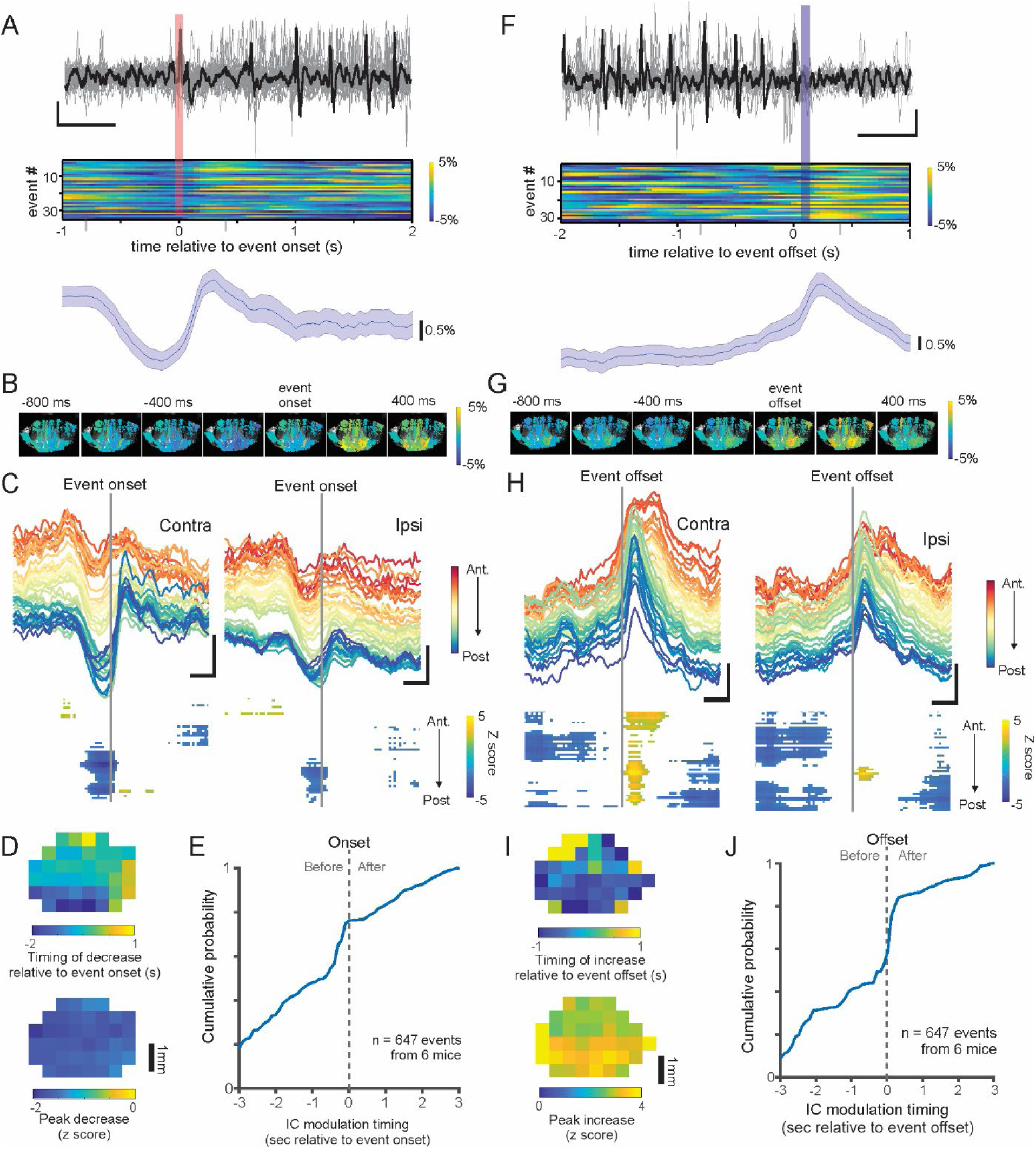
Widespread and early cerebellar changes also occur for hippocampal electrographic-only seizure events. A. Hippocampal LFP (stacked, one trace is shown in black for easy visualization; top) and calcium cerebellar signal (averaged across ICs) by seizure event (rows; middle heat plots) for all electrographic events recorded in an example recording day in an example mouse, aligned to event onset (vertical pink bar). Bottom trace: average (across events and ICs) calcium signal recorded from the cerebellum. Shading represents SEM. B. Parasagittal IC calcium changes, aligned to the timing of event onset, displaying a shorter time window, from 800ms prior to the first hippocampal ictal spike to 400ms after. Note the widespread depressed cerebellar calcium signals immediately prior to event onset. C. Top: Cerebellar calcium changes for this example animal and recording session averaged across events, by location within the cerebellum. Bottom: Corresponding significance-thresholded heat plots, indicating that significant decreases are observed prior to the onset of electrographic events, particularly in posterior regions. D. Average spatial maps of all ICs with significant decreases prior to electrographic event onset across all animals, indicating the timing of significant decreases is earliest for posterior ICs (top), but that the magnitude of the decrease is similar across the window (bottom). E. Cumulative probability of timing of significant modulation for all significantly modulated ICs, aligned to event onset. Note the inflection at ∼600 msec prior to hippocampal event onset. F-J, same as A-E but for electrographic event offset, illustrating widespread increases in cerebellar calcium activity at event offset. Scale bars: 0.5 s, 0.1mV A and F; 0.5 s, 2 z C and H.

For cerebellar modulation relative to IEDs, the ΔF/F of each IC was aligned to the peak of each IED event in the hippocampal LFP. For this analysis, only IEDs that were well isolated, i.e. more than 5 seconds from any other electrographic epileptiform event, were selected. For IEDs, the frequency of IEDs across recordings led to a limited, skewed, distribution of activity during interictal IED-free time periods. Thus, in order to determine significant IC modulation with respect to IEDs, we instead computed the average IC ΔF/F in a pre-IED window of 5-3 seconds prior to the IED to normalize and threshold the ΔF/F of each IC in the immediate peri-IED window. The peri-IED timing of IC modulation was computed for all ICs with significant modulation using a ±3 z-threshold. To visualize peri-IED changes, for each significantly modulated IC, the timing of the first significant change, as well as the average pre-IED modulation 500 msec before an IED and average post-IED modulation 500 msec after the IED were computed and averaged using the same bins as for behavioral and electrographic seizures to generate timing and modulation maps as in **Figure 6E**. In order to compare cerebellar IED-related changes across time, cerebellar IED-related changes were calculated by comparing the pre-IED ΔF/F (250ms before IED) and immediately post-IED ΔF/F (250ms after IED). For changes relative to behavioral seizures, sufficient high quality IEDs were present for 8 of 11 imaged behavioral seizures; IED dynamics relative to behavioral seizures draws from 8 events. Peri-IED ΔF/F changes were averaged across IEDs by 50 second bins relative to behavioral seizure onset (**Fig. 7A-B**) or by 2 day time bins relative to putative SUDEP event (**Fig. 7C-D**). Changes relative to behavioral seizure onset and/or putative SUDEP event were then examined using Pearsons’ correlations.

**Figure 6.**
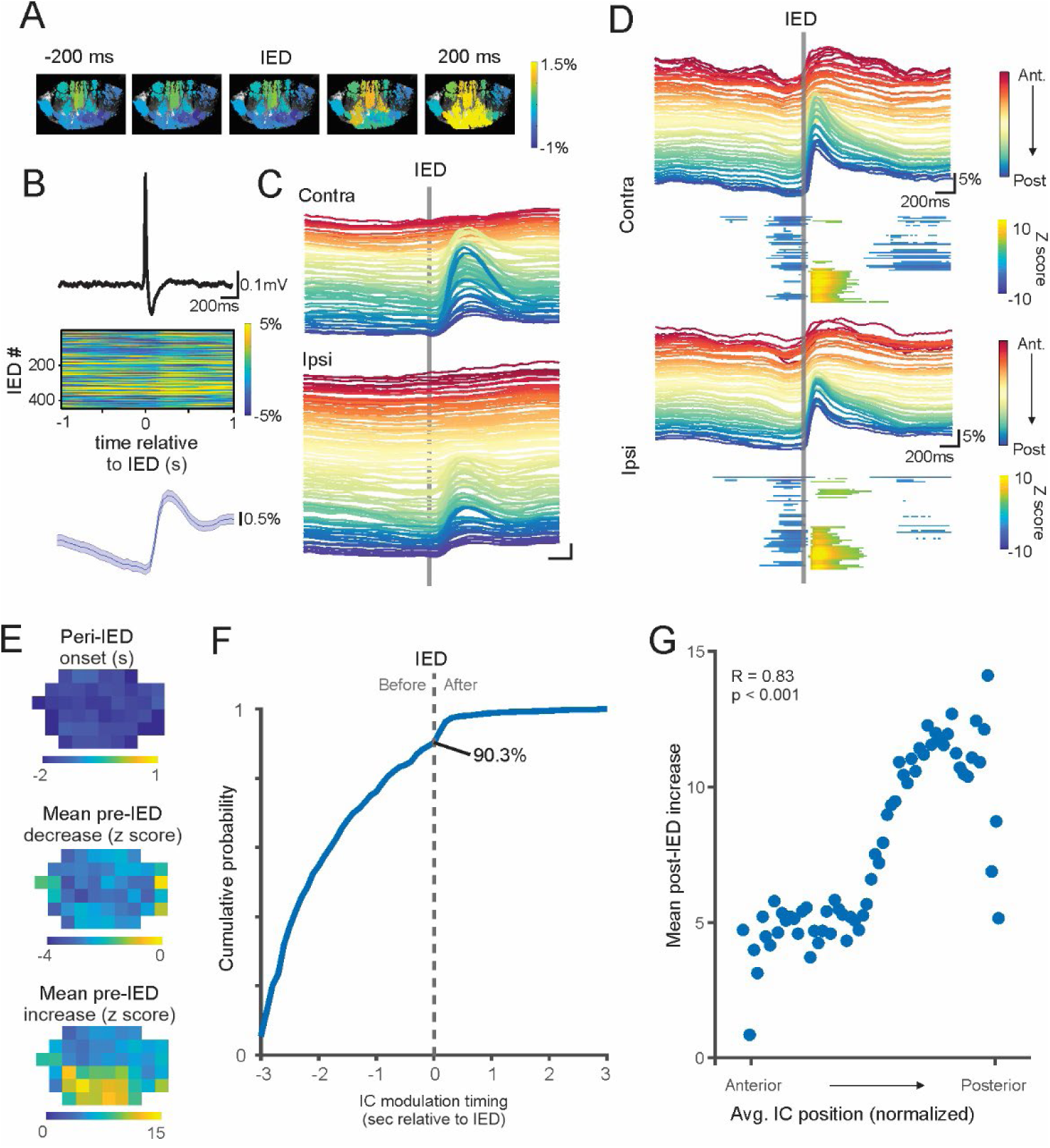
Hippocampal IEDs are associated with early and widespread cerebellar alterations, with cerebellar decreases immediately prior to hippocampal electrographic changes, followed by increased cerebellar calcium activity. A. Average fluorescence changes for parasagittal ICs aligned to IED onset in an example mouse on an example recording day, indicating decreased cerebellar activity prior to IED and a rebound in activation after the IED. B. Data from an example recording session in an example animal. Top: Average hippocampal LFP IED. Middle: Associated change in cerebellar calcium activity, averaged across all ICs, by IED. Bottom: Change in cerebellar activity averaged across IEDs. Shading represents SEM. Note that the decrease in average calcium activity begins >200 msec prior to any noticeable change in the hippocampal LFP. C. Cerebellar calcium changes for this example animal and recording session, averaged across IEDs, by location in the cerebellum. Scale bar: 200 msec, 1% ΔF/F. D. Contralateral (top) and ipsilateral (bottom) cerebellar calcium changes by anterior-posterior location in the cerebellum averaged across all IEDs recorded across all animals as ΔF/F signals (colored traces) and as thresholded activity (heat maps) across the same time window. Note the significant decreases in activation beginning prior to IED onset, and significant increases, particularly for posterior regions, after IED offset. E. Maps of average timing of significant modulation (top), mean pre-IED modulation (middle) and mean post-IED modulation (bottom) across all animals based on location within the cerebellar window, illustrating early decreases throughout the window, and strongest post-IED increases in the posterior regions. F. Cumulative plots of timing of significant modulation relative to IEDs for all significantly modulated ICs. G. Average post-IED increases based on anterior-posterior positioning in the cerebellar cortex (R = 0.83, p < 0.001, Pearson’s correlation).

**Figure 7.**
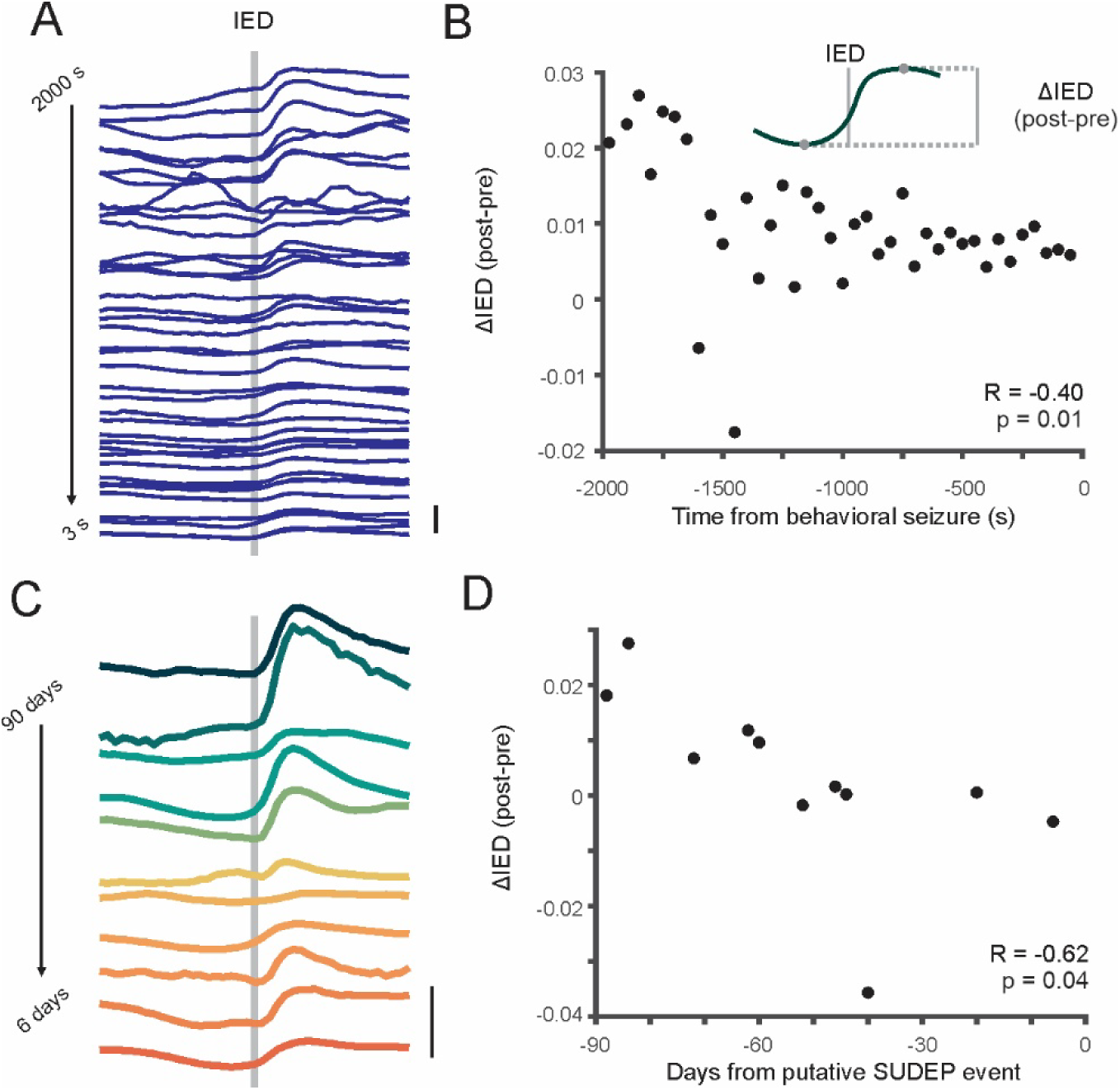
Dynamic cerebellar peri-IED activity is muted in the minutes leading up to a large behavioral seizure and in the days preceding a putative SUDEP event. A. Average peri-IED cerebellar response relative to behavioral seizure onset, beginning 2000 seconds prior to the seizure, in 50 second bins. B. Quantified peri-IED change (as indicated in schematic) for each line in A, illustrating more muted cerebellar response to IEDs as a behavioral seizure approaches (n = 8 behavioral seizures from 3 mice). C. Same as for A, but for an animal recorded over a three-month period prior to a putative SUDEP event. Traces illustrate the average peri-IED cerebellar response in 2 day bins. D. Same as for B, but for each trace in C. Scale bars A: 5% ΔF/F; C: 2% ΔF/F.

## Data availability

All data and original code generated for analysis will be made available upon reasonable request.

## Results

### Imaging during spontaneous behavioral seizures

To prevent direct damage to the cerebellum and to have a known site of injury, we utilized the unilateral intrahippocampal kainate mouse model (**Figure 1A**), in which the initial injury produces chronic epilepsy with hallmarks of unilateral temporal lobe epilepsy, including hippocampal sclerosis ^78^. Modified See-Shell ^74^ technology allowed chronic, widefield, monitoring of the cerebellar cortex ^75^, including lobules IV-VII, with concurrent local field potential (LFP) recordings in the hippocampus for seizure detection in awake, head-fixed, animals (**Figure 1B-D**). Due to the genetic approach taken, GCaMP6s was expressed sparsely in Purkinje cells across the entire cerebellum. This allowed both broad sampling of Purkinje cells across a wide area, and (due to the sparsity of expression) single cell resolution even with 1-photon imaging (**Figure 1D, inset**). Additionally, 1-photon imaging allowed simultaneous monitoring of both dendritic calcium signals (which are dominated by complex spikes ^98,99,103,104^) and somatic calcium signals in Purkinje cells ^75^. Unsupervised spatial independent component analysis provided unbiased segregation of the widefield Ca^2+^ signals into functional domains, which appeared as parasagittal bands and, separately, Purkinje cell somata (**Fig. 1E-F**)^75^.

A benefit of See-Shell technology is that it prevents deformation of the underlying tissue and allows repeated imaging of the same animal across many days. Repeated imaging allowed us to capture even relatively rare, spontaneous, overtly behavioral (minimum of Stage 3 on the Racine Scale ^105^) seizure events in chronically epileptic animals. Over 1,116 three-minute recording stacks, across 71 recording sessions from 6 animals, we successfully imaged the cerebellum during 11 spontaneous, recurrent, overtly behavioral, seizures (**Figures 2-4**).

Purkinje cell Ca^2+^ activity was clearly modulated by overtly behavioral temporal lobe seizures (**Figure 2, Supplementary Video 1**). In fact, seizures evoked widespread changes across the entire imaged cerebellum, including medial portions as well as lateral cerebellum (including Simplex and Crux I), both contralateral and ipsilateral to the seizure focus (**Fig. 2A & B maps**). This was apparent in both parasagittal domains (largely reflecting Purkinje cell dendritic calcium signals) (**Fig. 2, top panels**) and individual Purkinje cell somatic calcium signals (examples from different cerebellar regions illustrated in **Fig. 2** ΔF/F traces).

Notably, even before the onset of high amplitude hippocampal LFP ictal spikes, and even before any noticeable motor engagement, there was a marked, significant increase in Purkinje cell calcium signals (**Figure 2-3**). An increase in cerebellar calcium activity prior to overt behavioral seizure manifestations (**Fig. 2**) suggests cerebellar alterations are not simply due to motor engagement. An early increase in calcium signals was seen broadly across the cerebellum (medial, contralateral, and ipsilateral to the hippocampal LFP recording site) and in both the parasagittal and somatic domains (**Fig. 2**, **arrows**). An early increase was also visible across the anterior-posterior axis (**Fig. 3A-D**), with a very slight, but significant, earlier engagement in the posterior regions (significant IC modulation timing versus normalized anterior-posterior location of IC: R=-0.12, p=0.02). The majority of significantly modulated ICs, regardless of their location, exhibited the first significant increase in calcium signals ∼5-10 s prior to the onset of large hippocampal LFP ictal spiking (**Fig. 3E**).

As the seizures progressed, Purkinje cell hyperactivity gave way to marked hypoactivity (**Fig. 2, Supplementary Video 1**). This was apparent in both the parasagittal and somatic domains (**Fig. 2, Supplementary Figure 1**). Decreases in calcium activity occurred either at the onset of post-ictal depression (example in **Fig. 2A**, **Fig. 4A**) or prior to the offset of hippocampal LFP ictal spiking (examples in **Fig. 2B**, **Fig. 4B-C**). In either scenario, during periods of hippocampal LFP postictal depression, the cerebellum was essentially silent (**Fig. 2, Supplementary Video 1**).

Together, these data suggest that 1) the cerebellum undergoes widespread modulation during seizures, 2) the widespread modulation cannot be explained simply by the motor components of seizures, 3) that modulation of the cerebellum is not static, nor uniform, but rather goes through phases as a behavioral seizure progresses, and 4) hints that cerebellar modulation may even precede noticeable changes in the hippocampal LFP signal. However, as these are large behavioral seizures, potential caveats exist. We therefore turned to more subtle hippocampal epileptiform events.

### The cerebellum shows early and widespread engagement also during hippocampal electrographic-only seizures

The dorsal intrahippocampal kainate model of epilepsy, in addition to occasional overtly behavioral seizures, typically displays a higher frequency of seizure events which do not include motor manifestations ^79,80,86^ (**Supplemental Figure 2**). These electrographic-only seizure events (also referred to as paroxysmal discharges) allow for an examination of the cerebellum during epileptic events without the potential complication of motor engagement, and provide a tight temporal window to examine cerebellar alterations in relation to the hippocampal LFP.

We examined 647 hippocampal electrographic-only seizure events across 6 mice (**Figure 5**). As these events had variable durations, we aligned calcium signals to either the onset (first ictal spike) or the offset (last ictal spike) of the electrographic seizure event (**Fig. 5 A-E** and **Fig. 5 F-J**, respectively).

Remarkably, immediately (∼600ms) *prior* to electrographic seizure onset, calcium signals were decreased across the cerebellum (**Fig. 5A-E**). This early decrease was present in medial and lateral cerebellum (**Fig. 5B**), along the anterior-posterior axis (**Fig.5 C, D**), and both contralateral and ipsilateral to the hippocampal seizure focus (**Fig. 5C, D**). These changes were also visible in both the parasagittal domains and the calcium signals from Purkinje cell somata (**Supplementary Fig. 1**), and in both male and female animals (**Supplementary Figure 3**).

Across all ICs with significant modulation, >75% displayed significant modulation beginning prior to the first hippocampal-LFP ictal spike (**Fig. 5E**). Notably, as LFP recordings provide a faster readout than calcium imaging, the timing difference (i.e., cerebellar calcium changes *prior* to hippocampal LFP) cannot be explained by the differences in timescales associated with each recording modality.

Seizure offset showed an inverse profile, with a marked rebound in cerebellar signals as the seizure terminated (**Fig. 5F-J**). As with seizure onset, cerebellar changes with seizure offset were widespread, including medial, lateral, contralateral, ipsilateral, anterior, and posterior cerebellum (**Fig. 5F-I**), were present in both somatic and parasagittal components (**Supplemental Figure 1**), and in both male and female animals (**Supplemental Figure 3**).

### Interictal epileptiform discharges are also associated with widespread changes in the cerebellum, and changes in the cerebellum can precede hippocampal LFP changes

Interictal epileptiform discharges (IEDs) are very brief events, lasting on average ∼100ms in our recordings. Despite this brief duration, they were also associated with dramatic changes in calcium signals across the cerebellum (**Figure 6**). As with seizure events, cerebellar modulation peri-IED was bidirectional. Specifically, cerebellar calcium signals showed an initial decrease and then a rebound following the hippocampal IED. Astoundingly, even for IEDs, changes occurred in the cerebellum *prior* to any apparent changes in the hippocampal LFP (**Fig. 6 A-F**). The vast majority of modulated ICs showed significantly decreased activity in the seconds leading up to an IED, with approximately 90% of all IC modulation beginning *prior* to the IED (**Fig. 6F**). Again, as LFP recordings provide a faster readout than calcium imaging, the timing difference (i.e., cerebellar calcium changes *prior* to hippocampal LFP) cannot be explained by the differences in timescales associated with each recording modality, but could reflect the relative sensitivity of each modality. Post-IED increases were present in both parasagittal and somatic components (**Supplemental Figure 1**), and in both male and female animals (**Supplemental Figure 3**). IED rebounds were strongest in posterior (VI – VII) cerebellum (**Fig. 6E, G**).

Cerebellar alterations (specifically gray matter loss) are associated with an increased risk of Sudden Unexpected Death in Epilepsy (SUDEP)^45^. The occurrence of IEDs across many imaging sessions allowed us to examine peri-IED cerebellar dynamics across time. We specifically examined IED-related cerebellar changes with respect to two time points (**Figure 7**). First, we examined IEDs with respect to an approaching behavioral seizure (**Fig. 7A, B**). We found that the closer in time to a large behavioral seizure, the more muted the IED cerebellar response (**Fig. 7A, B**; peri-IED change in average cerebellar fluorescence versus time from behavioral seizures, R=-0.40, p = 0.01, Pearson’s correlation). Additionally, we had enough chronic recordings from one animal that eventually succumbed to a putative SUDEP event to examine cerebellar activity during IEDs in relationship to death. Here too we found a decrease in cerebellar IED responsiveness (**Fig. 7C, D**) in the days preceding the putative SUDEP event (peri-IED change in average cerebellar fluorescence versus time to death, R=-0.62, p = 0.04, Pearson’s correlation). Together, these results suggest that cerebellar IED responsiveness may serve as a measure of disease severity on both acute (minutes) and more extended (days-months) time scales.

Our results highlight distinct patterns of cerebellar modulation during three forms of spontaneous hippocampal epileptiform activity (summarized in schematic form in **Supplemental Figure 4**). Spontaneous, overtly behavioral seizures are associated with dramatic increases in cerebellar activity beginning several seconds prior to large amplitude spiking on the hippocampal LFP, and before observed behavioral changes. Conversely, electrographic only events are preceded by widespread decreases in cerebellar activation, mirrored by an increase in cerebellar activity at event offset. Interictal epileptiform events are also associated with a bidirectional cerebellar response, with a decrease prior to IED and a rebound immediately following. The magnitude of this bidirectional cerebellar response to IEDs was not static, and varied depending on the timing relative to a future behavioral seizure, and in the case of one animal, proximity to a putative SUDEP event.

## Discussion

Our results illustrate that widespread areas of the cerebellum (indeed, the entire imaging field of view -- encompassing anterior and more posterior cerebellum, and medial and lateral cerebellum) are modulated by large behavioral seizures, and *also* by seizures observable in the hippocampal LFP that were not accompanied by overt motor manifestations. Indeed, even brief hippocampal IEDs were accompanied by marked cerebellar alterations. Moreover, changes in the cerebellum were noted *prior* to changes in the hippocampal LFP. These results highlight that the cerebellum is not a passive bystander during hippocampal seizures, and should be considered as a potential key node in seizure networks.

While the cerebellum is often neglected in epilepsy research, a robust body of literature has indicated that it can be engaged by a variety of seizure types, including temporal lobe seizures ^30,31^. However, the extent of cerebellar modulation was unknown, and concerns about potential motor contributions lingered. As our results show robust cerebellar changes prior to motor engagement in seizures and widespread cerebellar changes for hippocampal electrographic-only seizures, cerebellar engagement in seizures does *not* simply reflect motor engagement.

Our results illustrate widespread alterations in the cerebellum during all ictal stages, including even *preictal* cerebellar modulation. Indeed, widespread and early changes were a universal finding across the different forms of epileptiform activity investigated. In this study, epilepsy was induced via unilateral intrahippocampal kainate injection. At the level of the hippocampus, electrographic-only seizures in this model of temporal lobe epilepsy typically remain unilateral only, with electrodes on the contralateral side often failing to record the event ^79^. Note that previous work has indicated that cerebellar electrodes *are* able to detect these seizure events ^33^ (indicating that cerebellar engagement is also notable with non-calcium imaging methods), but did not provide information regarding the extent of this engagement. The widespread and bilateral engagement of the cerebellum reported here, despite typical unilateral engagement of the hippocampus itself, is therefore particularly remarkable. While the anatomical pathways underlying the functional connectivity between the hippocampus and the cerebellum is an area of active investigation^20^, the cerebellum is connected via multi-synaptic and reciprocal loops to the neocortex in a crossed manner ^12,13^ (note that the cerebellum is ‘double-crossed’ with respect to the body). Therefore, cerebellar engagement contralateral to the seizure focus, rather than bilateral cerebellar engagement, may have been predicted. Bilateral engagement in the cerebellum during hippocampal electrographic-only seizures suggests bilateral functional connectivity between the hippocampus and the cerebellum. Prior work has also shown bilateral functional connectivity in the opposite direction (i.e., from the cerebellum to the hippocampus); cerebellar modulation ipsilateral or contralateral to the hippocampal seizure focus is able to inhibit hippocampal seizures ^33,44^. Again, the anatomical substrate underlying this bilateral functional connectivity is unclear, but may be mediated in part via cerebellar commissural connections, and has been shown to involve fastigial neurons projecting to the central lateral thalamus ^39^. Cerebellar engagement during hippocampal seizures was seen not only bilaterally, but also along the entire anterior-posterior extent of the imaging window, including vermal lobules IV/V - VII, as well as Simplex and Crus 1. The extent of cerebellar modulation was remarkable.

Additionally, these were spontaneous epileptiform events, observed in awake, behaving, animals. Changes in cerebellar activity were therefore observed above the ‘noise’ of ongoing cerebellar modulation with respect to behavior. Such marked cerebellar modulation during even relatively subtle electrographic-only seizure events and IEDs is therefore particularly striking.

While widespread and early changes were a universal finding across the different forms of epileptiform activity investigated, there were some notable differences as well (key patterns of cerebellar changes summarized in **Supplemental Figure 4**). In large, overtly behavioral seizures, we noted early ictal increases in Purkinje cell calcium signals along the anterior-posterior and medio-lateral axes, in both parasagittal domains (i.e. dendritic calcium signals) and in calcium signals in individual Purkinje cell somata. This early increase continued during the early ictal event, but as the seizure progressed, gave way to a marked decrease, with decreases often starting in more anterior portions and progressing posteriorly. At or prior to the onset of postictal depression in the hippocampal LFP, the cerebellum showed profound quiescence across the entire imaging field of view. In contrast, electrographic seizures which did not progress to large, overtly behavioral seizures, and which were uniformly of the “hypersynchronous” ^80^ type, showed a preictal *decrease* in cerebellar calcium signals, and a large postictal rebound, suggesting fundamental differences in cerebellar activity patterns across these two types of seizure events. Somewhat parallel to hippocampal electrographic-only seizure events, cerebellar calcium signals were decreased prior to hippocampal IEDs, followed by a large rebound in cerebellar activity.

The postictal rebound following electrographic seizures may reflect a release of inhibition as less severe seizures end, or -- especially given previous work showing that stimulating the cerebellum can inhibit seizures ^31,33,38,44^ -- potentially a sign of native seizure-suppressive mechanisms at play, although the impact on cerebellar nuclei remains to be determined. The profound cerebellar depression seen with more severe seizures may reflect a failure of native seizure-suppressive mechanisms, or simply extensive Purkinje cell inhibition. Such profound inhibition of the cerebellum may have consequences for postictal recovery and even SUDEP^45^. Note that the cerebellum projects to brainstem regions important for respiratory control, including the periaqueductal gray (PAG) and the parabrachial nucleus ^39,106,107^. In our study, the large number of IEDs, recorded over many imaging sessions, allowed an examination of how cerebellar responses to epileptiform activity may change with time and disease severity. We found that cerebellar responsiveness to IEDs was muted in the minutes leading up to a behavioral seizure, and anecdotally, in the days leading up to a putative SUDEP event. As noted above, cerebellar structural changes may be predictive of future SUDEP events in human epilepsy patients ^45^.

Importantly, sustained hippocampal ictal activity was not required to observe changes in cerebellar calcium signals, as even brief IEDs were associated with widespread cerebellar changes. This included decreased calcium signals immediately *prior* to the IED, followed by increased calcium signals, especially in posterior regions. Neocortical changes have also been noted for IEDs ^108^, illustrating their potential for widespread effects. Therefore, despite the cerebellum’s historic relative neglect in epilepsy, it may not be entirely surprising that activity in the cerebellum is also altered during IEDs. However, the timing of the cerebellar changes with regard to IEDs is particularly surprising. The reported modulation of the neocortex by hippocampal IEDs is in keeping with hippocampal IEDs inducing neocortical changes, with neocortical changes *lagging* hippocampal IEDs ^108^. Even calcium imaging from the hippocampus itself does not indicate hippocampal changes *prior* to the IED; calcium imaging from the contralateral hippocampus during IEDs illustrates changes during and after, rather than changes *prior* to the IED ^109^. Specifically, an increase in hippocampal inhibitory interneuron activity *during* hippocampal IEDs resulting in a decrease in hippocampal pyramidal cell activity is reported ^109^. Our data illustrating that changes in the cerebellum can actually *precede* the hippocampal IED is therefore particularly remarkable, and may question the order of cause-and-effect. Alternatively, the cerebellum may be particularly sensitive to hippocampal changes, such that even early hippocampal changes -- which are not yet discernable in the LFP ^71^ nor noted via CA1 calcium imaging ^109^ -- are able to cause marked, widespread, cerebellar changes. In considering relative timing, it is also important to consider the location of the IED/seizure focus. In this study, epilepsy was induced via a direct insult to the hippocampus, and the hippocampus is the presumed seizure focus. However, it is possible that the hippocampal electrode still reflects IED propagation. In this scenario, early cerebellar changes may be paired with the true IED onset, rather than being immediately prior to the IED. However, this would require faster transmission of the IEDs to the cerebellum than to neighboring hippocampus. In short, while there are multiple potential explanations for how changes could occur in the cerebellum prior to hippocampal LFP changes, any imagined scenario is remarkable and requires serious reconsideration of how the cerebellum is viewed in relationship to epileptiform activity.

Decreased cerebellar activity was also noted prior to hippocampal electrographic seizures. While widespread changes immediately prior to an IED are particularly remarkable, preictal changes are also noteworthy. Preictal changes are not a finding unique to the cerebellum or this model of epilepsy, and may reflect early altered activity or a seizure prone state. Preictal changes have been noted in superior colliculus activity prior to seizures in a rat model of absence epilepsy^110^, and preictal changes have been observed in neocortical interneurons in heat-induced seizures prior to seizure onset ^62,63^. Specifically, a preictal increase in activity in neocortical somatostatin-expressing neurons, and a deficit in parvalbumin-interneuron synchronization, has been observed with heating in a mouse model of Dravet syndrome ^62,63^. A preictal appearance of superhubs in pentylenetetrazol-treated zebrafish has also been noted^53^, as well as preictal changes in glial calcium signals during copper-or acute kainate-induced seizures ^111,112^.

While our work is the first work to perform calcium imaging in the cerebellum during spontaneous epileptiform activity, previous work has imaged the cerebellum during chemically evoked seizures in zebrafish ^67–69^. Ictal increases in zebrafish cerebellum were also noted ^69^, suggesting cerebellar engagement during seizures is not limited to rodent epilepsy.

Additionally, while calcium imaging cannot be performed in patient populations, other imaging modalities (including SPECT and MRI) have illustrated cerebellar changes in epilepsy, including cerebellar ictal hyperperfusion in TLE ^35,113–116^, which in some patients can even be more obvious than in the presumed seizure focus itself ^116^. Similarly, in keeping with the large rebound we find in cerebellar calcium signals immediately following a hippocampal IED, prior work using EEG-fMRI in temporal lobe epilepsy patients found an increase in blood-oxygen-level dependent (BOLD) responses in areas including the cerebellar cortex^117^. The temporal resolution of BOLD imaging may have prevented detection of smaller pre-IED decreases in cerebellar activity. In addition to their significance for epilepsy and seizure networks, our results provide further support for the robustness of cerebellar-hippocampal interactions ^20^, and highlight a need to consider the cerebellum in cognitive and “extra motor” domains, as the cerebellum is frequently excluded from cognitive neuroimaging studies ^118^.

Additionally, while our work has focused on the cerebellum during hippocampal epileptiform events in a model of temporal lobe epilepsy, the cerebellum may be more widely relevant to the epilepsies^30^. Changes in the cerebellum during IEDs have also been noted in other patient populations and studies^119^, including frontal lobe epilepsy patients^117^. Absence seizures are associated with both cerebellar BOLD increases in human patients^120^ and altered Purkinje cell firing in animal models^32,36^. LFP and unit recordings have illustrated cerebellar impacts across a range of animal seizure models (reviewed in reference ^30^).

The cerebellum is rarely believed to be the seizure focus, though a large number of case reports illustrate that the cerebellum can be ictogenic ^30,121–124^. A range of semiologies have been noted in patients with cerebellar lesions or tumors, including hemifacial seizures, progressive myoclonia, and tonic-clonic seizures ^121–124^. The cerebellar lesion and presumed focus has been associated with altered metabolism and perfusion on PET and SPECT images, scalp EEG recordings can indicate the cerebellum as the maximum source of aberrant activity, and intraoperative depth recordings have illustrated epileptiform discharges from the lesion ^49,50,125^.

Moreover, these cerebellar epileptiform discharges are reported to *precede* intracranial EEG abnormalities in the neocortex ^50^. Confirming the ictogenic locus, removal of the cerebellar lesion can produce seizure freedom ^30,50,125^.

Except in such cases where there is an apparent lesion, the cerebellum is rarely included in a surgical workup for epilepsy patients. Increased monitoring may lead to increased observations of the cerebellum being engaged during epileptiform activity in clinical populations. Indeed, in rare clinical depth recordings from the cerebellum in patients without noted cerebellar lesions, cerebellar engagement has been observed ^34^. This includes a patient with chronic epilepsy following encephalitis 7 years prior, where strong and *early* electrographic engagement was recorded from the fastigial electrode ^34^, further underscoring the clinical relevance of our findings. Notably, our work utilized the intrahippocampal kainate model of temporal lobe epilepsy, avoiding direct insult to the cerebellum. We still found widespread (and early) engagement of the cerebellum in epileptiform events. It may therefore be beneficial to include the cerebellum as an area of interest even in cases where the seizure focus is believed to be well-defined.

We show here that the cerebellum shows early and widespread engagement during diverse forms of epileptiform activity. Previous work has shown that the cerebellum can also be a target for seizure control. As the cerebellum can both modulate and be modulated by epileptiform activity, the cerebellum should be considered as an important node in seizure networks.

## Competing interests

The authors declare no financial competing interests.

**Supplemental Video.** Widefield cerebellar imaging stack captured during the spontaneous behavioral seizure in **Figure 2A**. Top left panel is the hippocampal LFP recorded during the entire stack, bottom left panel is a zoomed in view of the real time hippocampal LFP aligned to the imaging stack. Note the increased, bilateral activation of the cerebellum beginning several seconds prior to the high amplitude spiking of the behavioral seizure, the sustained posterior activation during the seizure, and the depression beginning in anterior regions of the cerebellum prior to seizure offset. Bright postictal flashes are also apparent with concurrent postictal spikes in the hippocampal LFP.

---

**Supplemental Figure 1.**
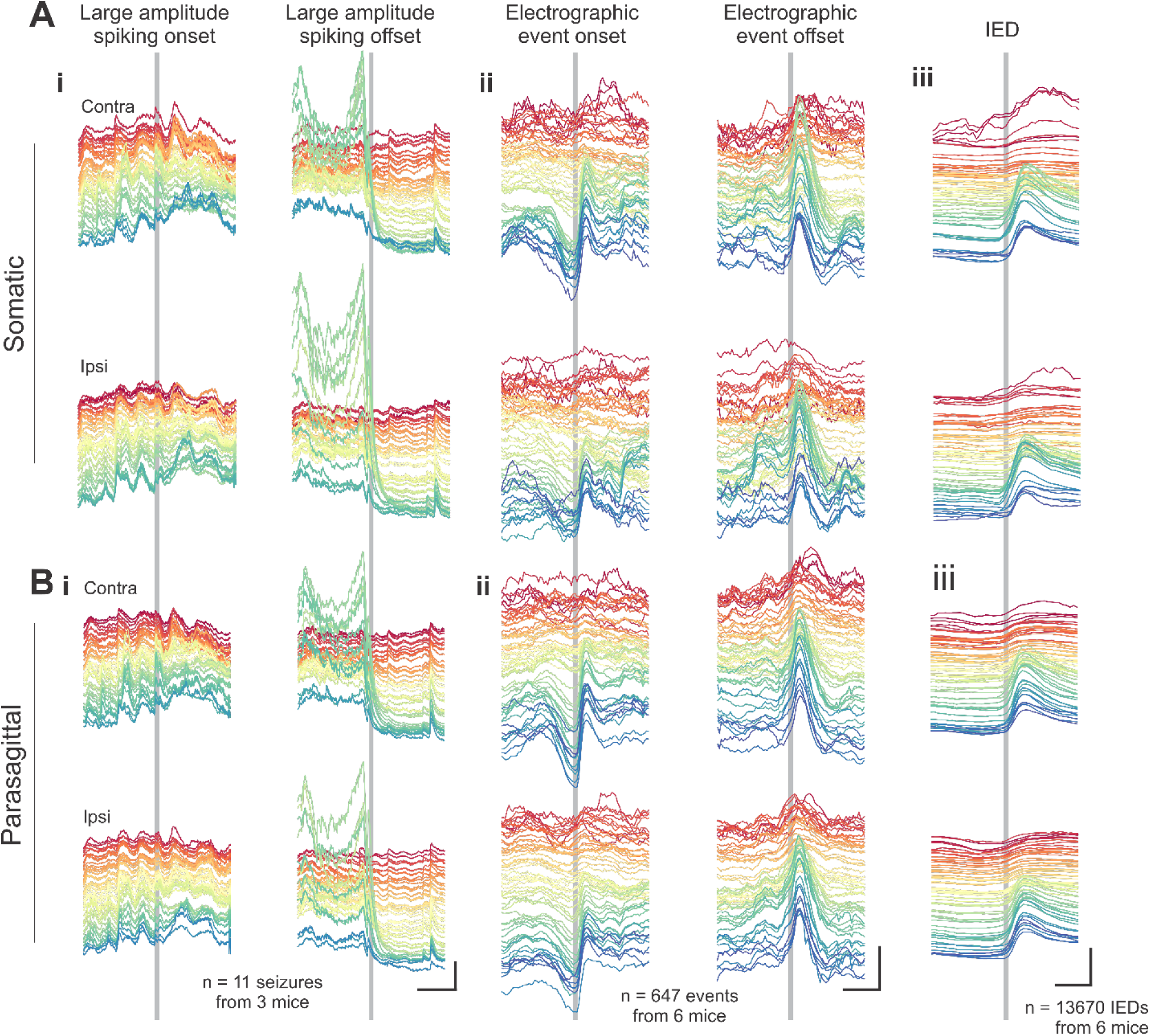
Noted changes are observable in both parasagittal and somatic components of cerebellar calcium signals. Average calcium signals by cerebellar area (warm colors indicate more anterior cerebellum) for contralateral (top) or ipsilateral (bottom) somatic (A) or parasagittal (B) domains for seizures with an overt behavioral component (i), electrographic seizures without an overt behavioral component (ii), or IEDs (iii). Scale bars: i: 20% change in ΔF/F, 5s; ii: 2%, 1s; iii: 2%, 0.5s.

**Supplemental Figure 2.**
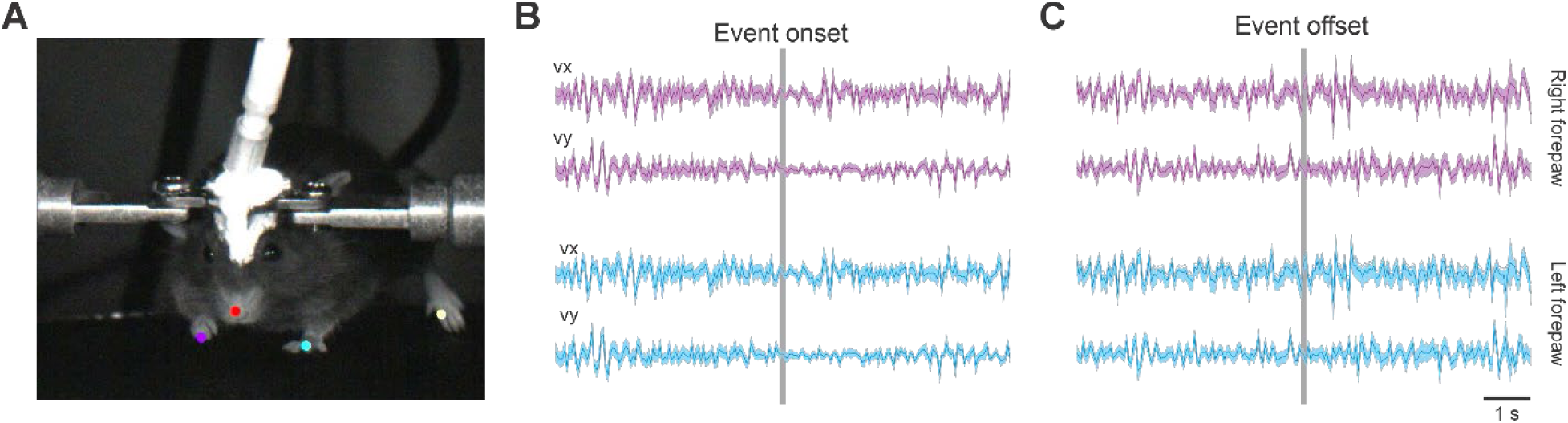
Hippocampal electrographic-only seizure events are not accompanied by observed motor manifestations. A. DeepLabCut ^91^ markerless tracking allowed automated tracking of the right (purple) and left (blue) forepaws during imaging sessions. B-C. Electrographic seizures were not associated with any notable changes in the movement of either forepaw, at seizure onset (B) nor seizure offset (C). Shown is relative position for each paw in the video image for the x (vx, top) or y (vy, bottom) axes. Shading represents SEM. Data represents paw location data for 244 electrographic seizure events from 6 animals. Scale bar: 1s.

**Supplemental Figure 3.**
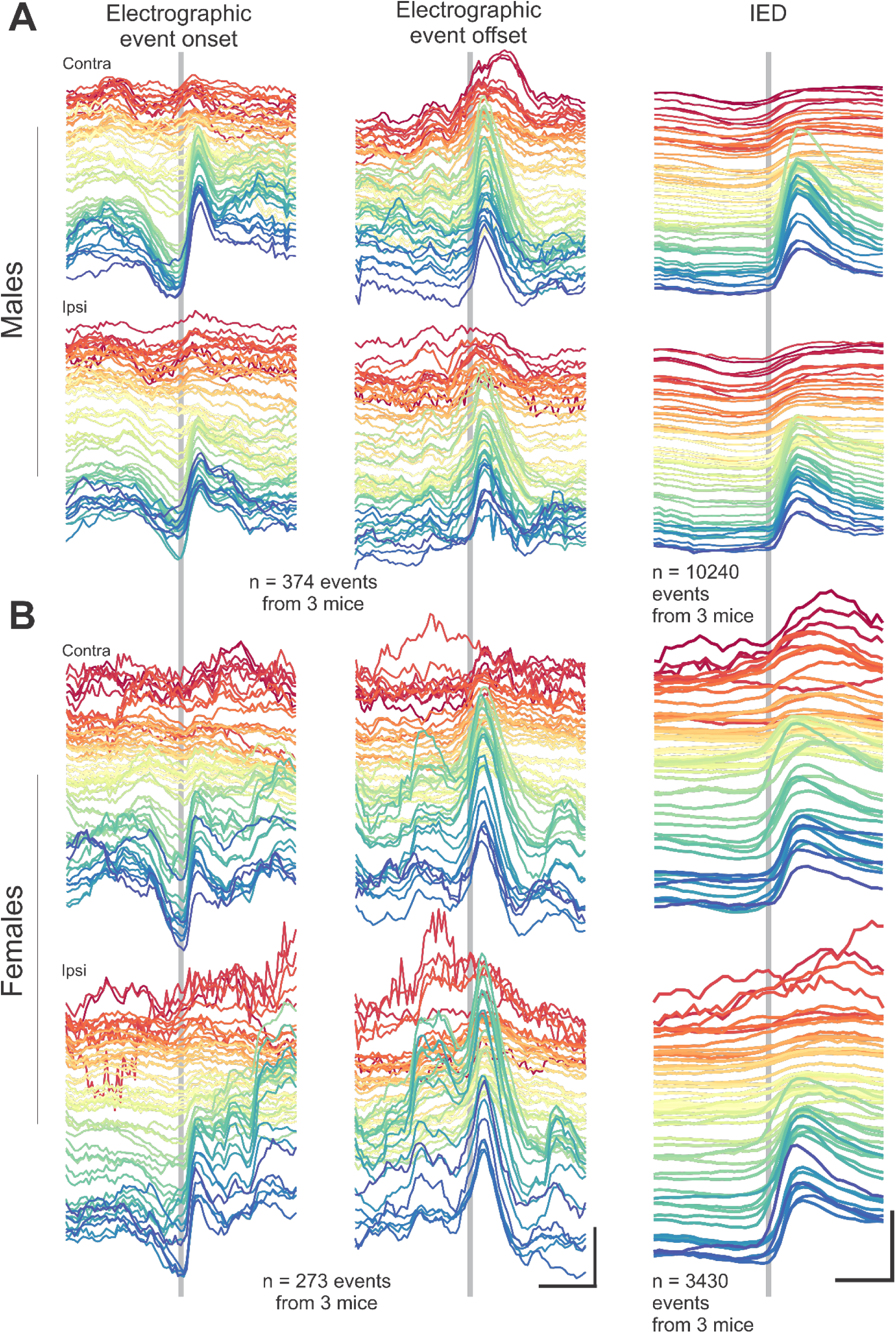
Noted cerebellar changes during electrographic seizures and IEDs are present in both male and female animals. Average cerebellar responses by cerebellar region (top: contralateral, bottom: ipsilateral cerebellum; warmer colors represent more anterior regions) for male (A) and female (B) animals aligned to electrographic seizure onset (left), offset (middle), or IED peak (right). Note that the pre-ictal decrease, the post-ictal rebound, and the initial slight-dip followed by large post-IED rebound is visible for both male and female animals. Scale bars: 1 second, 2% ΔF/F.

**Supplemental Figure 4.**
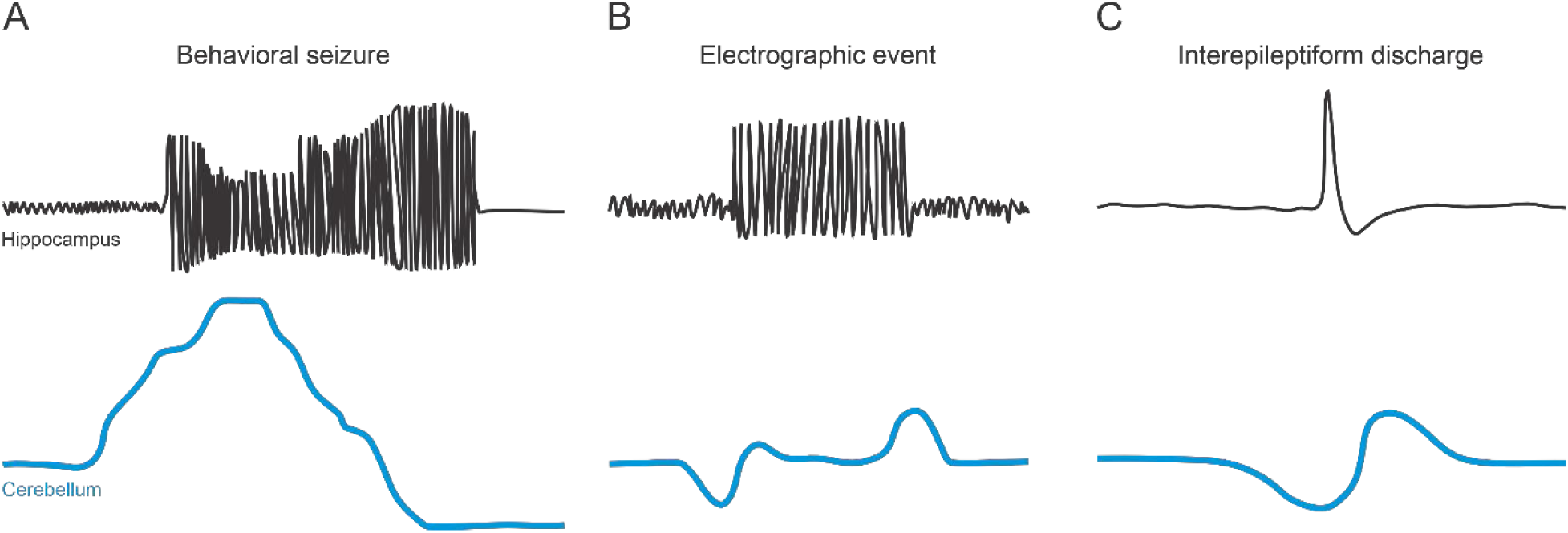
Summary schematic of findings. We find widespread cerebellar engagement during hippocampal epileptiform activity that begins prior to obvious changes in the hippocampal LFP. During behavioral seizures (A), robust increases in cerebellar activation are observed, which begin several seconds prior to large amplitude hippocampal spiking. Dramatic decreases in cerebellar engagement often begin several seconds prior to the offset of high amplitude spiking, and persist throughout the postictal period. During electrographic-only events (B), decreased cerebellar activation begins prior to event onset, with an increase at event offset. Interictal epileptiform discharges (IEDs) are associated with a decrease in cerebellar activity beginning prior to LFP changes followed by a rebound thereafter. We found that the features of this bimodal response to IED could be associated with upcoming behavioral seizures and, in one animal, death due to a putative SUDEP event.

